# The effects of running shoe stack height on running style and stability during level running at different running speeds

**DOI:** 10.1101/2024.11.19.624278

**Authors:** Cagla Kettner, Bernd Stetter, Thorsten Stein

## Abstract

The footwear market contains a wide variety of running shoe solutions aiming at optimizing performance and minimizing injuries. Stack height is one of the most highly discussed design features of running shoes, but its effects are not yet well understood. This study investigated the effects of different shoes differing mainly in their stack heights (High: 50 mm, Medium: 35 mm & Low: 27 mm) on running style and stability during treadmill running at 10 and 15 km/h. A total of 17 healthy experienced runners participated in this study. The kinematic data were recorded with a 3D motion capturing system. The running style was investigated by a dual-axis framework with duty factor (DF) and leg length normalized to step frequency (SFnorm). Additionally, the ratio of landing to take-off duration, the lower body joint angle time series in the sagittal and frontal planes, the vertical center of mass oscillation (COMosc), and the stiffness parameters (kver & kleg) were compared for different conditions. The stability was analyzed using linear (i.e. discrete frontal ankle parameters) and nonlinear methods (i.e. Maximum Lyapunov Exponent for local dynamic stability of head, trunk, hip, and foot, and detrended fluctuation analysis of stride time). High resulted in longer ground contact relative to stride time (DF) compared to Low. The higher the stack height, the higher was the COMosc. Furthermore, High led to a longer foot eversion during stance compared to Medium. In addition, the local dynamic stability of the hip decreased with High in comparison with Low. The higher stack heights (≥ 35 mm) led to a lower SFnorm at 15 km/h but not at 10 km/h. The remaining shoe effects were independent of the running speed. Findings showed that changes in stack height can affect running style. Furthermore, the highest stack height resulted in changes related with instabilities (i.e., longer foot eversion and lower hip dynamic stability) which may be a critical issue in terms of injuries and performance. However, this study did not include joint load analysis or running performance measures such as VO2. Future studies may benefit from the combination of analysis approaches to better understand stack height effects on running injuries and performance.

## 1 Introduction

The footwear market is flooded with running shoe solutions that combine various features, such as midsole stiffness, shape of toe box, stack height and heel-to-toe drop. Ultimately, running shoes should be optimized to help improve running performance as well as avoid injuries (Hoitz et al., 2020; Mai et al., 2023). Despite a high number of studies, there is still an ongoing debate on what effects different shoe features have on running performance and injuries (Burns & Tam, 2020; Willwacher & Weir, 2023). On this basis, it is crucial to evaluate the effects of different shoe features in detail to understand their biomechanical function (Mai et al., 2023).

Stack height, or the thickness of the sole, is a highly discussed feature of running shoes, especially since it is limited by the new World Athletics regulation (World Athletic Council, 2022). According to the World Athletic Shoe Regulations (approved on the 22.12. 2021 and effective from 01.01.2022), the thickness of a shoe sole cannot be greater than 40 mm for none-spikes shoes. The measurements are done at the center of the forefoot and heel, which are located at 12% and 75% of the internal length of a shoe, respectively. This regulation was introduced to avoid the possible performance benefits by use of advanced footwear technologies (Burns & Tam, 2020; Ruiz-Alias et al., 2023a). However, the actual effects of stack height on running performance and injuries are not yet well understood (Barrons et al., 2023a, 2023b; Hoogkamer, 2020; Ruiz-Alias et al., 2023a; Ruiz-Alias et al., 2023b; Willwacher & Weir, 2023). Various studies reported that increased stack height may lead to performance increases, particularly due to elongated effective leg length (e.g., Burns & Tam, 2020; Ruiz-Alias et al., 2023a). An increase in effective leg length can lead to an increase in stride length, which may ultimately help to increase running speed and finish a race faster (Burns & Tam, 2020). However, there are also studies suggesting that stack height changes alone cannot be sufficient to increase running performance (Bertschy et al., 2023). Furthermore, the added mass due to a higher stack height may also impair performance by decreasing economy (Burns & Tam, 2020; Hoogkamer, 2020). Besides the running economy perspective, higher stack heights may also decrease ankle frontal plane stability (e.g., Hoogkamer, 2020; Ruiz-Alias et al., 2023a), which may be a critical issue in terms of running performance and injuries. With respect to stack height, optimum levels possibly exist for certain conditions (e.g., running speed, stiffness of the ground material etc.). However, there is still no clear consensus on where the optimum is and how to determine it (Hoogkamer, 2020). One reason for this may be that the combined effects of individual features are difficult to decompose (Burns & Tam, 2020). For example, even if only stack height is increased, the amount of material increases, and inevitably the mass also increases. Another challenge is the non-standardized measurement and reporting protocols both with respect to the shoes (e.g., measurement of the stack height or reporting the shoe features) and study design (e.g., self-selected speed instead of a fixed one) that make the results difficult to compare (Frederick, 2020; Hannigan & Pollard, 2020).

Running style, defined as the “visually distinguishable movement pattern of a runner” (van Oeveren et al., 2021) was suggested to be important in terms of running performance as well as injuries (Barnes & Kilding, 2015; Floría et al., 2024; Folland et al., 2017; Mann et al., 2015; Nijs et al., 2022; Saunders et al., 2004). In a synthesis paper on the biomechanics of running and running styles, step frequency normalized to leg length (SFnorm) and duty factor (DF) were proposed as sufficient to provide an overview of a running style (van Oeveren et al., 2021). However, while spatiotemporal characteristics provide fundamental information on running style, they may still not be enough to understand it completely. For example, TenBroek et al. (2014) focused on various discrete joint angle variables and showed that running style changed when wearing shoes with different stack heights. In addition, a recent study (Koegel et al., 2024) aimed to investigate individual biomechanical running responses and clustered runners into three groups with distinct running patterns. Their study showed that leg stiffness (kleg) and vertical center of mass oscillation (COMosc) contributed most to the clustering. This finding underlined the distinguishing characteristic of these variables (Koegel et al., 2024). Additionally, a decreased COMosc was associated with a better running economy (Folland et al., 2017). The studies focusing mainly on the effects of shoe stack height during running reported that a higher stack height may increase the effective leg length, therefore increase the step length as well as stance time (Burns & Tam, 2020; TenBroek et al., 2014); but a comprehensive analysis with respect to running style has not yet been conducted. Furthermore, while stiffness and center of mass (COM) movement changes due to different shoes have been analyzed in a few studies (Chambon et al., 2014; Kulmala et al., 2018), none focused on stack height, especially on the higher height (i.e. ≥ 35 mm). Further data here may help to improve understanding of stack height effects. More concretely, Kulmala et al. (2018) compared two different running shoes which differed in multiple features and showed that high cushioning shoes increase impact loading especially at a higher speed. Chambon et al. (2014) analyzed running shoes with stack heights only up to 16 mm and found increased stance time with increasing stack height but no effects on ground reaction force or tibial acceleration. Consequently, there still exists a research gap in the study of higher stack heights in relation to running style.

In addition to running style, running stability is a commonly discussed issue, as it is crucial in terms of running performance as well as injuries (Barrons et al., 2023a; Bruijn et al., 2013; Frank et al., 2019; Hoenig et al., 2019; Promsri et al., 2024; Schütte et al., 2018), especially with respect to stack height (Barrons et al., 2023a; Esculier et al., 2015). In dynamic systems, stability is defined as the ability to compensate for perturbations (Strogatz, 2015). In the context of running, stability can be defined as the ability to maintain functional running movement (e.g., without falls) despite the presence of perturbations (e.g., fatigue or different shoes (Frank et al., 2019; Schütte et al., 2018)). Stability can be subdivided into orbital, local and global stability in the context of running (Dingwell & Kang, 2007), although several studies overlooked this distinguishment. A gait pattern can be locally unstable but still maintain orbital or global stability (Riva et al., 2013; Santuz et al., 2020a).

Maximum Lyapunov Exponent (MLE), which is a nonlinear method, has often been applied to operationalize the local dynamic stability of body regions or joint angles in running studies (Winter et al., 2024). It was shown that expertise level, running speed and fatigue affect local stability (Frank et al., 2019; Hoenig et al., 2019; Mehdizadeh et al., 2014). Another study on the effects of the shoe on stability showed that minimalist shoes did not differ from traditional shoes, but higher stack heights were not investigated (Frank et al., 2019). Another approach is to analyze the stability of a more global variable (i.e., stride interval) by looking into their long-range correlations using detrended fluctuation analysis (DFA) (Agresta et al., 2019; Hausdorff et al., 1996). The rationale behind this approach is that the inherent variability in gait data is not a random fluctuation but is a part of long-range power-law correlations. From a practical point of view, it means that the fluctuations in strides are related to variations in much earlier strides (Hausdorff et al., 1996). An increase in these long-range correlations (e.g., due to manipulations in footwear or foot strike patterns (Fuller et al., 2016)) is interpreted as a decreased ability to adapt running strides to a changed external condition, and therefore a reduced global stability (Agresta et al., 2019; Jordan et al., 2006).

Besides MLE and DFA, linear approaches have been used in various studies to operationalize the stability - particularly based on the movement of the ankle joint in the frontal plane (Barrons et al., 2023a; Isherwood et al., 2021). Thereby, discrete variables such as peak eversion/inversion foot angle or total eversion duration during stance are used to operationalize ankle stability or to assess the risk of running-related injuries (Barrons et al., 2023a, 2024; Becker et al., 2017, 2018; Hannigan & Pollard, 2020; Isherwood et al., 2021; Kuhman et al., 2016; TenBroek et al., 2014). For example, Barrons and colleagues (2023a) reported a higher peak eversion angle for recreational runners with a higher stack height (45 mm vs. 35 mm), which was interpreted as a lower local stability of ankle Since the studies analyzing stability mostly focus on one analysis approach (e.g., either linear (Law et al., 2019) or nonlinear analysis (Frank et al., 2019)), it becomes difficult to build a comprehensive understanding which combines results from different analysis approaches.

Addressing the above-mentioned research gaps, this study investigated the effects of shoes with different stack heights on running style and stability during level running at different speeds. It was hypothesized that the shoes with higher stack heights (≥ 35 mm) lead to changes in running style (H1) and reduced stability (H2). Additionally, it was expected that a higher running speed (i.e. 15 km/h) leads to more pronounced changes compared to a lower speed condition (i.e. 10 km/h) (H3).

## 2 Methods

### 2.1 Participants

Seventeen experienced healthy male runners participated in this study (age: 25.7 ± 3.9 years, height: 1.77 ± 0.04 m, mass: 68.1 ± 6.0 kg, shoe size: EU 42-43, running activity per week: 4.2 ± 1.8 days and 33.7 ± 22.4 km). All participants provided written informed consent prior to the measurements. The study was approved by the ethics committee of the Karlsruhe Institute of Technology (KIT).

### 2.2 Experimental protocol

The measurements were conducted on a treadmill (h/p/cosmos Saturn, Nussdorf-Traunstein, Germany). The experimental protocol began with a warm-up and familiarization to the treadmill and three slope conditions (0%, −10% and 10%) by running at self-selected speed with their own shoes for 5 min in total (Paquette et al., 2024). This study focuses only on the 0% condition. Then, the measurement blocks were repeated for three running shoes with different stack heights tested in this study (High: 50 mm, Medium: 35 mm and Low: 27 mm, measured at the heel, US men size 9) in a parallelized order (Table 1 & Fig. 1). High and Medium were made out of the same materials and included curved carbon infused rods (Barrons et al. 2023a) but Low had only TORSIONRODS to increase force transfer (Adidas Adizero RC4). For each shoe condition, the protocol began with a familiarization to the current shoes by running at self-selected speed for 3 min and walking at 5 km/h for 2 min in the level condition (0%) (Paquette et al., 2024). Afterwards, a total of six running trials with different slope and speed conditions were performed. For each slope condition, first the slow than the fast running condition was performed. For the 0% condition, 10 km/h and 15 km/h were chosen based on the test measurements and previous studies (Fadillioglu et al., 2022; Kulmala et al., 2018; Selvakumar et al., 2023). Each of the six running trials lasted 90 s. Before each trial the participants ran for 25-40 s until the treadmill reached the target speed. There were 1 min walking breaks between the slow and fast running conditions and a 2 min standing break between the slope conditions to prevent exhaustion. A Borg scale (Borg, 1998) was used before each running trial to control the exhaustion level (i.e. the next trial did not begin unless the Borg score was <12 on a 6-20 scale). After each pair of shoes, a visual analogue scaled comfort questionnaire for the stability, cushioning and comfort in a range of 0-100% (from low to high) was collected (Fadillioglu et al., 2024).

**Figure 1:**
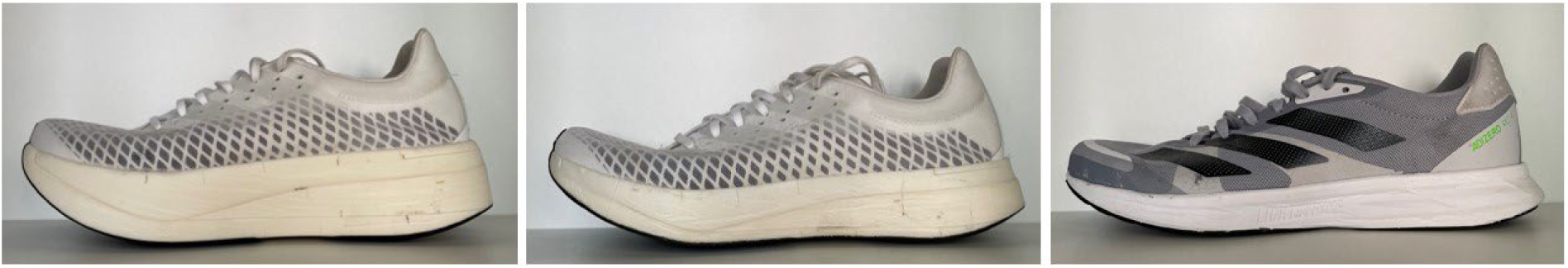
The tested shoes. On the left side High, in the middle Medium and on the right side Low.

**Table 1:** Features of the tested running shoes.

| Shoe | Mass<br>(g) | Forefoot height<br>(mm) | Heel height<br>(mm) | Heel-to-toe drop<br>(mm) |
| --- | --- | --- | --- | --- |
| **High** | 268 | 43 mm | 50 mm | 7 mm |
| **Medium** | 220 | 28 mm | 35 mm | 7 mm |
| **Low** | 219 | 19 mm | 27 mm | 8 mm |

### 2.3 Data acquisition, data processing and biomechanical modelling

A 3D motion capture system (Vicon Motion Systems; Oxford Metrics Group, Oxford, UK; 200 Hz) with 65 reflective markers was used to capture full body kinematics. Raw kinematic data were processed offline with Vicon Nexus V2.12 to reconstruct the 3D coordinates of the markers. Further data processing was performed in MATLAB (2023a, MathWorks Inc., Natick, USA). Marker data were filtered using a low pass Butterworth filter (4^th^ order, cut-off frequency 10 Hz (Gullstrand et al., 2009)). A full-body model (modified version of the OpenSim Hamner Running Model (Delp et al., 2007; Hamner et al., 2010)) was used to calculate joint kinematics using the OpenSim. The model was iteratively scaled for each subject separately until the maximum marker error was less than 2 cm and the root mean square error less than 1 cm. The weights for all markers were equal. The initial contact events were detected based on the local minimum in velocity curves of the mean of heel and toe markers (Leitch et al., 2011) and the toe-off events based on the maximum extension in sagittal knee angles (Fellin et al., 2010). The parameters for linear and nonlinear analyses were determined for 20 and 100 running strides, respectively, of the left side (Ogaya et al., 2021; Riazati et al., 2019). For the linear analysis, the 20 strides were selected from the first ∼15-35% of the 90 s recording period to minimize the number of missing markers during labelling of data, as some markers fell off due to the dynamic characteristic of running or sweating towards the end of the trials. For the nonlinear analysis, the 100 strides were selected from the ∼5-85% of the 90 s recording period.

### 2.4 Data analysis

#### 2.4.1 Running style

The SF_norm_ was calculated based on initial contacts, and normalized to leg length (l_0_) (Eq. 1) (Hof, 1996; van Oeveren et al. 2021).

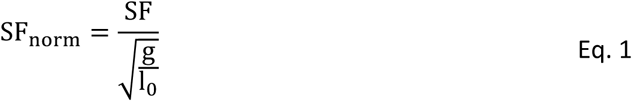

The DF was calculated as the ratio of stance time to the twice the sum of stance and flight time (Eq. 2) (Fadillioglu et al., 2022; van Oeveren et al., 2021).

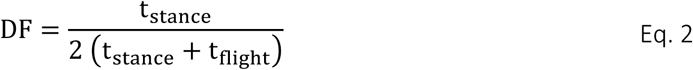

The results of SF_norm_ and DF are based on the dual-axis framework for running style (van Oeveren et al., 2021).

The braking and propulsion timings during stance were estimated using the maximum knee flexion event (Ciacci et al., 2010). The ratio of braking to propulsion duration (L2T) was calculated (Eq. 3) to estimate the braking to propulsion asymmetries (da Rosa et al., 2019).

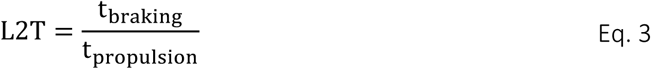

Frontal and sagittal angle time series of the ankle, knee and hip joints were time-normalized to stance phase (101 points) for each cycle separately using a cubic spline interpolation (Möhler et al., 2022). The individual time series were compared with each other across different shoe and speed conditions.

COM_osc_ was calculated for the stance phase by the range of motion of COM in vertical direction. Vertical stiffness (k_ver_) and leg stiffness (k_leg_) were estimated using Eq. 4 and Eq. 5, respectively (Garofolini et al., 2022; Morin et al., 2005).

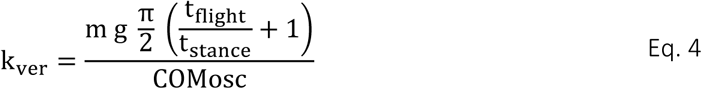

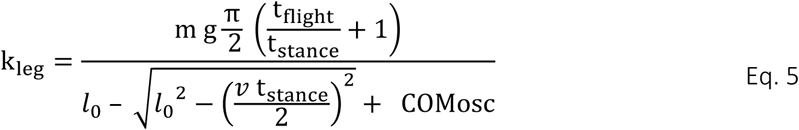

#### 2.4.2 Running stability

In the linear analysis, the following were calculated: total time at eversion normalized to stance time (t_eversion_), maximum eversion (MAX_eversion_) and maximum inversion (MAX_inversion_) angles during stance phase, and ankle frontal angle at initial contact (IC_inversion_) (Barrons et al., 2023a, 2024; Hannigan & Pollard, 2020; Isherwood et al., 2021; TenBroek et al., 2014).

In the nonlinear analysis, the local dynamic stability of foot, hip, trunk and head in the vertical axis were determined using MLE of marker clusters (each consisting of four markers) attached to the corresponding body regions (Ekizos et al., 2018; Hunter et al., 2023; Jordan et al., 2009; Look et al., 2013; Winter et al., 2024). Time series data of 100 strides were normalized to 10,000 points (100 strides x 100 points) (Hoenig et al., 2019a; Raffalt et al., 2019). Embedding dimension (*m*) was determined based on the false nearest neighbor method for all trials (Stergiou, 2016; Wallot & Mønster, 2018). The largest *m* over all trials was used for each region (*m_foot_* = 10, *m_hip_* = 8, *m_trunk_* = 10, *m_head_* = 8) (Hoenig et al., 2019a). Time lag (*τ*) for the reconstruction of the space was determined based on the first local minimum of the average mutual information curves (Stergiou, 2016; Wallot & Mønster, 2018). The median *τ* over all trials for each body region was used for further calculations (*τ_foot_* = 26, *τ_hip_* = 23, *τ_trunk_* = 23, *τ_head_* = 23) (Hoenig et al., 2019a). Rosenstein’s algorithm was used to estimate the divergence curve (Eq. 6) (Ekizos et al., 2017; Rosenstein et al., 1993). The slope of short-term phase (λ) was used to calculate MLE (Mehdizadeh et al., 2014). A lower λ indicated a higher dynamic local stability, and vice versa.

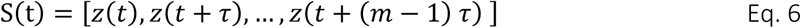

A DFA of stride time was performed to assess long-range correlations of running stride (Hunter et al., 2023; Jordan & Newell, 2008; Mann et al., 2015). First, the data series *x* was shifted by its average, summed cumulatively to obtained the integrated data (Eq. 7) and segmented into non-overlapping windows with a size of Δn varying between 4 and 24 time points. In the second step, integrated data were locally fit to a line by the least square method in each window, and average fluctuation of the time series around the estimated trend was calculated (Eq. 8). The second step was repeated for all the window sizes Δn (Bryce & Sprague, 2012).

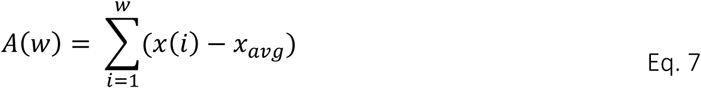

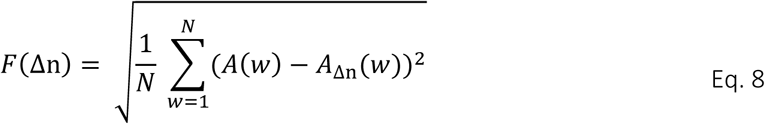

If the data show power law scaling characteristics, then a logarithmic (i.e., log-log) plot of F(Δn) vs. Δn is expected to be linear (Eq. 9). The slope of this curve gives the scaling component DFA-α which is used to quantify the long-range correlations of data. If the DFA-α lies between 0.5 and 1, the data show a self-similarity characteristic (e.g., a short stride is more likely to follow a short stride). In this range, the larger the DFA-α, the higher the persistence of data, which is interpreted as reduced stability (Agresta et al., 2019; Jordan et al., 2009).

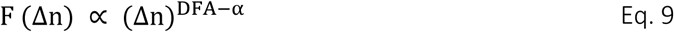

### 2.5 Statistics

#### 2.5.1 Running style

The statistical tests for discrete parameters were performed in SPSS (Version 29.0, SPSS Inc., IBM, Armonk, NY, USA). The average of the 20 cycles in each running trial was calculated for comparisons. The normality and sphericity were checked by the Kolmogorov-Smilnov and Mauchly tests, respectively. Greenhouse-Geisser estimates were used to correct for violations of sphericity. Repeated measurements analyses of variance (rmANOVAs) were performed to compare conditions where the independent factors were the shoe (High, Medium, Low) and speed (10 km/h, 15 km/h). Eta squared (η^2^) was calculated to estimate the sizes of the main effects (small η^2^ ≤ 0.06; medium 0.06 < η^2^ < 0.14; large 0.14 ≤ η^2^). Paired t-tests with Bonferroni-Holm corrections were calculated as *post-hoc* tests. In the case of a global shoe effect, the mean data at 10 km/h and 15 km/h were used in *post-hoc* tests, whereas in the case of a significant interaction effect, the *post-hoc* tests were performed for 10 km/h and 15 km/h separately. Cohen’s d was calculated to estimate the effect sizes for *post-hoc* tests (small d ≤ 0.50; medium 0.50 < d < 0.80; large 0.80 ≤ d) (Cohen, 1988). The significance level was set *a priori* to α = 0.05.

The angle time series within each joint dimension were compared using the statistical parametric mapping (SPM) toolbox in MATLAB (spm1d toolbox) (Pataky et al., 2019). Normality was checked using the normality tests provided in the toolbox. In the case of non-normality, nonparametric alternatives were conducted with 1,000 iterations. In the case of significant effects, the paired t-tests were calculated as *post-hoc* tests. The significance level was set *a priori* to α = 0.05.

#### 2.5.2 Running stability

The statistical tests were also performed in SPSS. For the comparisons of linear analysis, the average of the 20 cycles in each running trial was calculated. Within the nonlinear analysis, a single value was obtained for 100 cycles of each running trial. The Kolmogorov-Smilnov and Mauchly tests were used to check the normality and sphericity, respectively. rmANOVAs were performed to compare conditions with the independent factors shoe (High, Medium, Low) and speed (10 km/h, 15 km/h). In the case of a global shoe effect, paired t-tests with Bonferroni-Holm corrections were performed as *post-hoc* tests in which the mean data of 10 km/h and 15 km/h were used to compare shoe conditions independent of the speed effects. η^2^ and d were calculated to estimate the effects sizes as explained above. The significance level was set *a priori* to α = 0.05.

## 3 Results

### 3.1 Running style

The discrete parameters used for running style analysis are represented in Table 2. Fig. 2 visualizes the results for SF_norm_ and DF in the dual-axis framework. The SFnorm showed significant interaction between shoes and speed (p = 0.001). The *post-hoc* analysis showed that the SF_norm_ of Low was significantly higher than High (p = 0.046) and Medium (p = 0.012); and that of High was higher compared to Medium (p = 0.038) at 15 km/h.

**Figure 2:**
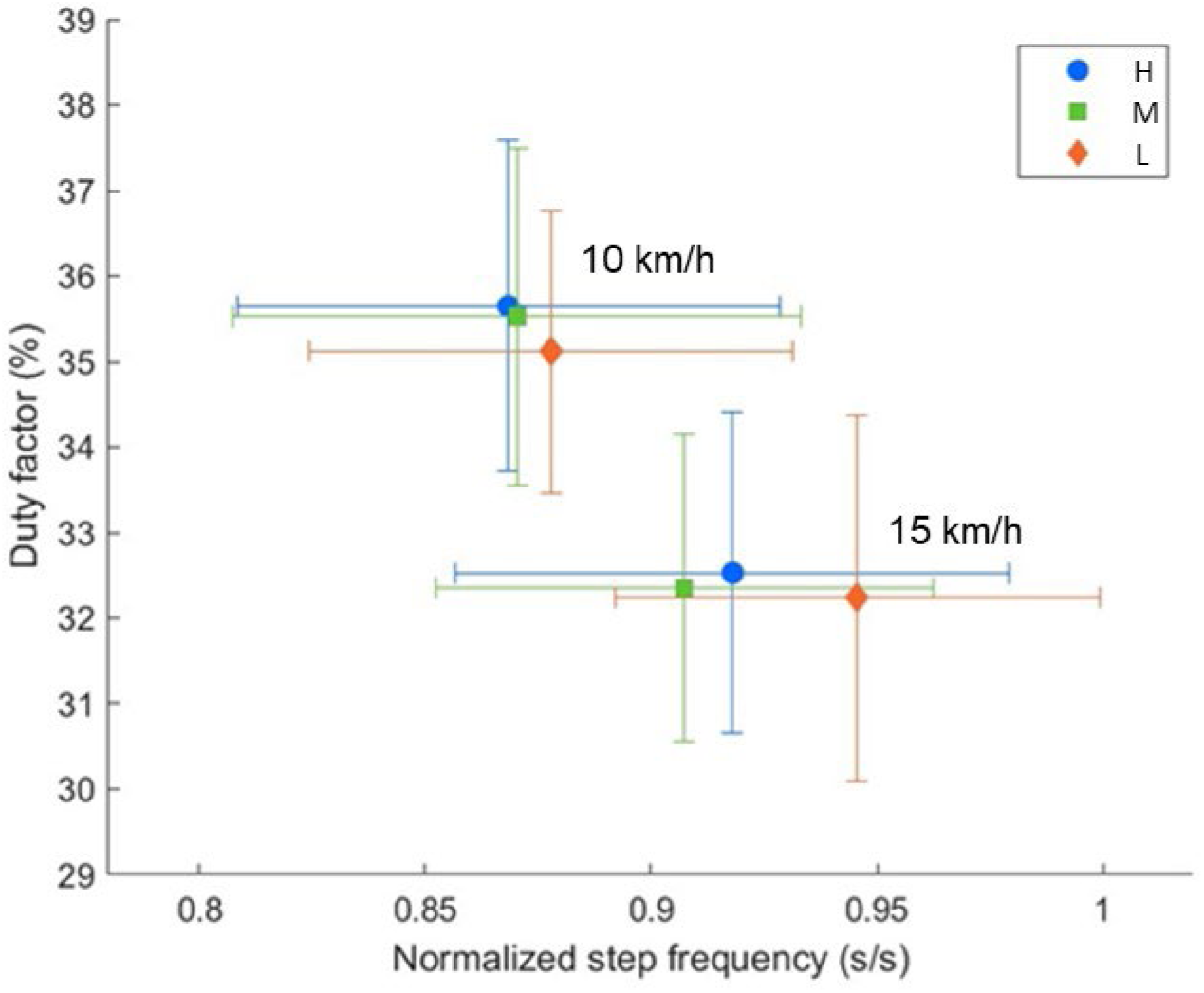
Running style analysis based on the dual-axis framework (van Oeveren et al., 2021) for different shoes (stack height, H: 50 mm, M: 35 mm & L: 27 mm) and running speeds (10 km/h and 15 km/h). The symbols indicate the mean values of each shoe and the error bars show standard deviations.

**Table 2:** The discrete parameters used for running style analysis: step frequency normalized to leg length (SF_norm_), duty factor (DF), ratio of landing to take-off duration (L2T), vertical center of mass oscillation (COM_osc_), vertical stiffnes (k_ver_) and leg stiffness (k_leg_). Shoe stack heights were abbreviated as H: 50 mm, M: 35 mm & L: 27 mm. Descriptive statistics is given as mean ± standard deviation. Unless otherwise specified, the *post-hoc* results were calculated for mean data at 10 km/h and 15 km/h. Significant results are highlighted in bold.

| $SF_{norm}$ (s/s) | | | | | | | | |
| --- | --- | --- | --- | --- | --- | --- | --- | --- |
| | 10 km/h | 15 km/h | rmANOVA | p | $\eta^2$ | *Post-hoc*<br>10 km/h | p | \|d\| |
| H | 0.87 ± 0.06 | 0.92 ± 0.06 | Shoe | 0.112 | 0.146 | H vs. M | 0.342 | 0.10 |
| M | 0.87 ± 0.06 | 0.90 ± 0.05 | Speed | **<0.001** | **0.727** | H vs. L | 0.735 | 0.17 |
| L | 0.88 ± 0.05 | 0.94 ± 0.05 | Interaction | **0.001** | **0.348** | M vs. L | 0.576 | 0.14 |
|  |  |  |  |  |  | *Post-hoc*<br>15 km/h | p | \|d\| |
|  |  |  |  |  |  | H vs. M | **0.038** | **0.46** |
|  |  |  |  |  |  | H vs. L | **0.046** | **0.53** |
|  |  |  |  |  |  | M vs. L | **0.012** | **0.73** |
| DF |  |  |  |  |  |  |  |  |
| | 10 km/h | 15 km/h | rmANOVA | p | $\eta^2$ | *Post-hoc* | p | \|d\| |
| H | 0.36 ± 0.02 | 0.32 ± 0.02 | Shoe | **0.018** | **0.222** | H vs. M | 0.176 | 0.23 |
| M | 0.35 ± 0.02 | 0.32 ± 0.02 | Speed | **<0.001** | **0.862** | H vs. L | **0.021** | **0.67** |
| L | 0.35 ± 0.01 | 0.32 ± 0.02 | Interaction | 0.510 | 0.041 | M vs. L | 0.144 | 0.37 |
| L2T (s/s) |  |  |  |  |  |  |  |  |
| | 10 km/h | 15 km/h | rmANOVA | p | $\eta^2$ | | | |
| H | 0.64 ± 0.06 | 0.67 ± 0.05 | Shoe | 0.952 | 0.003 | | | |
| M | 0.64 ± 0.05 | 0.67 ± 0.04 | Speed | 0.107 | 0.164 | | | |
| L | 0.64 ± 0.06 | 0.66 ± 0.07 | Interaction | 0.494 | 0.046 | | | |
| $COM_{osc}$ (m) | | | | | | | | |
| | 10 km/h | 15 km/h | rmANOVA | p | $\eta^2$ | *Post-hoc* | p | \|d\| |
| H | 0.084 ± 0.012 | 0.075 ± 0.011 | Shoe | **0.013** | **0.287** | H vs. M | **0.035** | **0.47** |
| M | 0.082 ± 0.012 | 0.072 ± 0.010 | Speed | **0.001** | **0.698** | H vs. L | **0.012** | **0.74** |
| L | 0.081 ± 0.011 | 0.070 ± 0.001 | Interaction | 0.925 | 0.005 | M vs. L | **0.030** | **0.58** |
| $k_{ver}$ (kN/m) | | | | | | | | |
| | 10 km/h | 15 km/h | rmANOVA | p | $\eta^2$ | *Post-hoc* | p | \|d\| |
| H | 35.66 ± 6.48 | 44.53 ± 7.15 | Shoe | **0.014** | **0.282** | H vs. M | 0.060 | 0.40 |
| M | 36.69 ± 6.01 | 45.86 ± 8.26 | Speed | **<0.001** | **0.886** | H vs. L | **0.009** | **0.75** |
| L | 37.57 ± 7.45 | 46.92 ± 7.45 | Interaction | 0.938 | 0.004 | M vs. L | **0.010** | **0.71** |
| $k_{leg}$ (kN/m) | | | | | | | | |
| | 10 km/h | 15 km/h | rmANOVA | p | $\eta^2$ | | | |
| H | 19.29 ± 3.49 | 17.42 ± 2.95 | Shoe | 0.154 | 0.110 | | | |
| M | 19.41 ± 2.96 | 17.03 ± 2.26 | Speed | **<0.001** | **0.576** | | | |
| L | 20.28 ± 3.82 | 18.01 ± 2.63 | Interaction | 0.719 | 0.020 | | | |

The DF was significantly different between shoes (p = 0.018). According to the *post-hoc* analysis, the DF was higher for High compared to Low independent of speed (p = 0.021). The L2T showed no significant effects (Table 2).

The ankle, knee and hip joint time series are shown in Fig. 3 and Fig. 4 for the sagittal and frontal planes, respectively. The SPM analysis revealed significant shoe effects for the sagittal and frontal ankle angles. The hip and knee angles did not differ significantly between the shoes. No interaction effects were detected. *Post-hoc* SPM t-tests revealed that Medium led to significantly higher inversion compared to High and Low (both p < 0.001). *Post-hoc* results in the sagittal plane did not reach the significance level.

**Figure 3:**
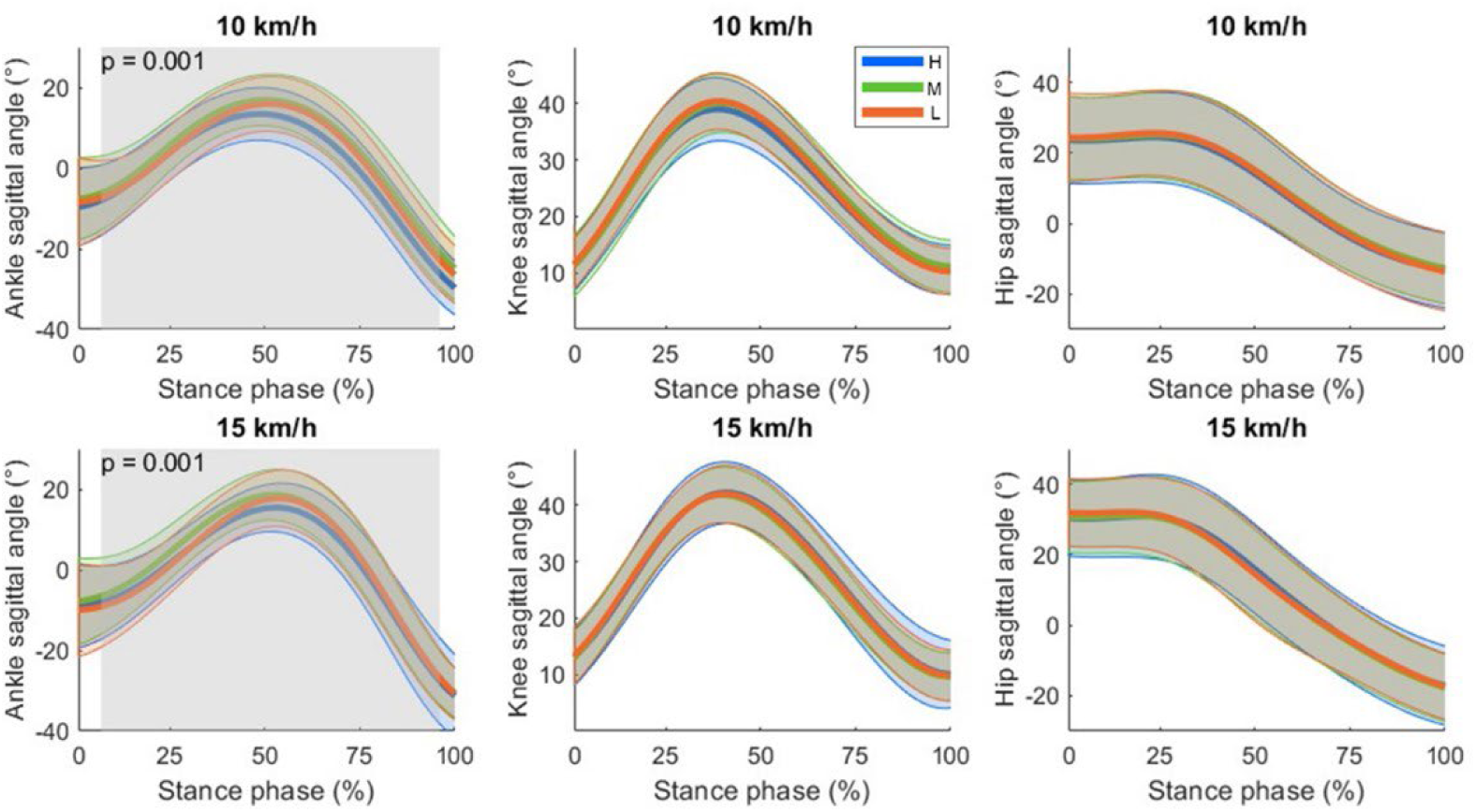
Sagittal plane joint angle time series for ankle, knee and hip as mean (thicker lines) ± standard deviations (upper and lower thinner lines). By convention, a positive angle indicated a flexion for all the joints (dorsiflexion in case of the ankle). Significant shoe differences independent of the running speed are highlighted with shaded areas. The corresponding p-values are also displayed. Shoe stack heights were abbreviated as H: 50 mm, M: 35 mm & L: 27 mm.

**Figure 4:**
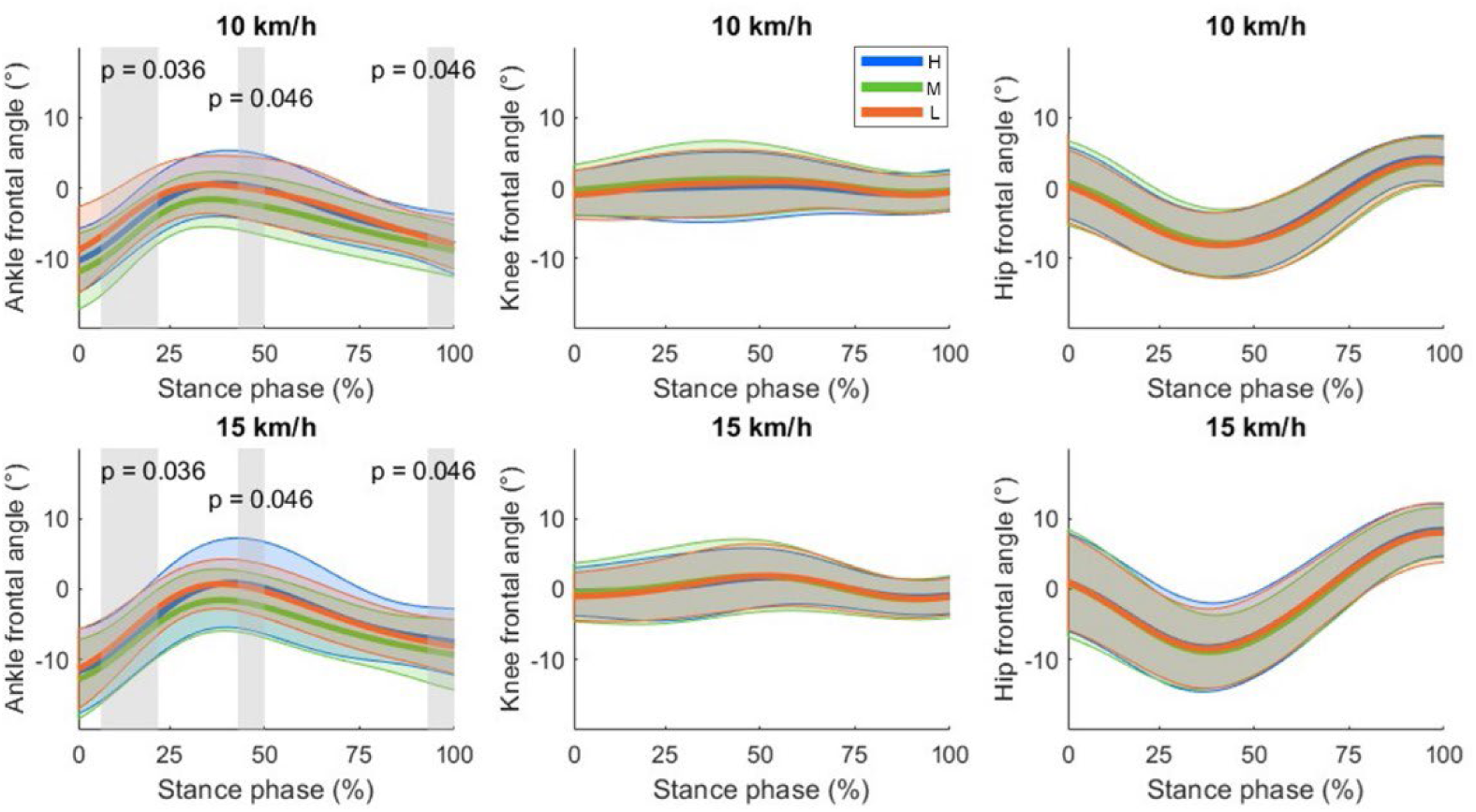
Frontal plane joint angle time series for ankle, knee and hip as mean (thicker lines) ± standard deviations (upper and lower thinner lines). By convention, a positive angle indicated an abduction for the knee and hip, and an eversion for the ankle. Significant shoe differences independent of the running speed are highlighted with shaded areas. The corresponding p-values are also displayed. Shoe stack heights were abbreviated as H: 50 mm, M: 35 mm & L: 27 mm.

The COM_osc_, k_ver_ and k_leg_ results are shown in Table 2 and Fig. 5. The COM_osc_ differed significantly between the shoes (p = 0.013). The *post-hoc* tests revealed that all pairwise comparisons between shoes were significant (Table 2). The k_ver_ showed significant shoe effects (p = 0.014). The *post-hoc* tests revealed that Low had the highest k_ver_.

**Figure 5:**
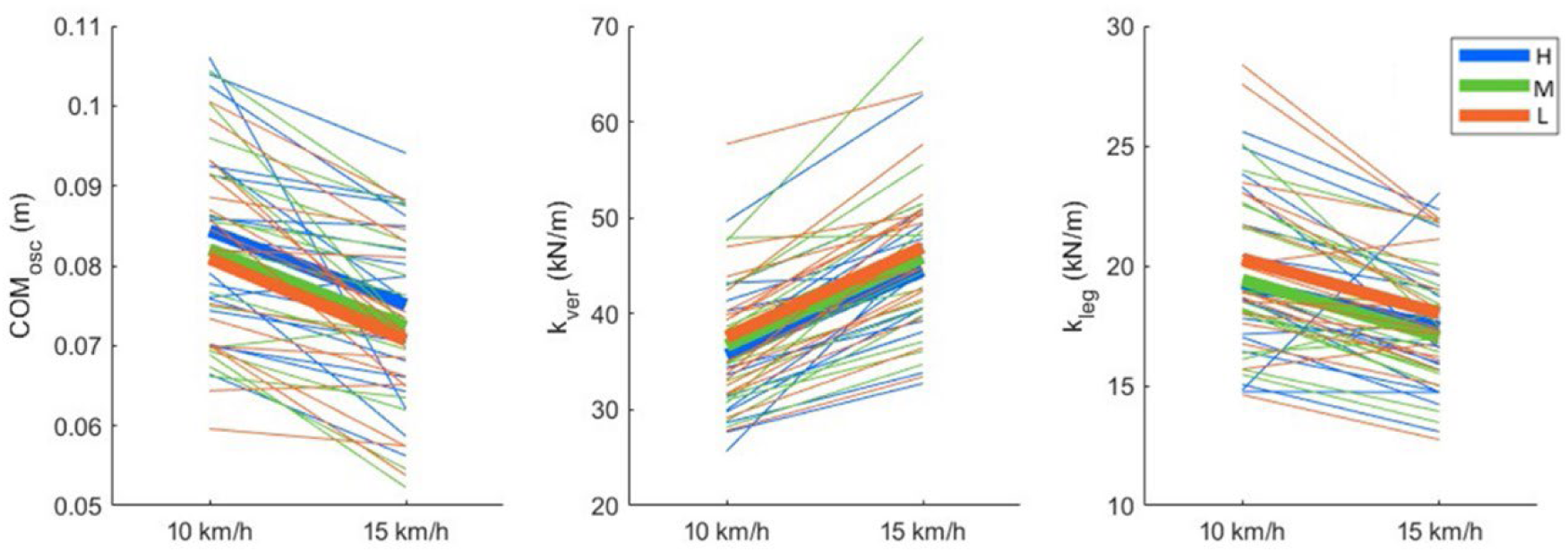
Vertical oscillation of the center of mass (COM_osc_), vertical stiffness (k_ver_) and leg stiffness (k_leg_) results. Thicker lines show mean values for each shoe, whereas thinner lines show each participant separately. Shoe stack heights were abbreviated as H: 50 mm, M: 35 mm & L: 27 mm.

### 3.2 Running stability

#### 3.2.1 Linear analysis

The results are summarized in Table 3. The t_eversion_ differed between the shoes (p = 0.035). The *post-hoc* tests revealed that Medium spent less time in eversion compared with High (p = 0.006) and Low (p = 0.046) independent of the running speed (Table 3). MAX_eversion_ showed significant effects for shoes (p = 0.006). The *post-hoc* tests showed that MAX_eversion_ during stance was lower in Medium compared to both High (p = 0.002) and Low (p = 0.010). The results of MAX_inversion_ showed significant differences for shoes (p = 0.005). According to the *post-hoc* tests, the MAX_inversion_ during stance was lower (higher in amount) in Medium compared to both High (p = 0.002) and Low (p = 0.004).

**Table 3:** The parameters used in the linear analysis of running stability: total time at eversion normalized to stance time (t_eversion_), maximum eversion (MAX_eversion_) and maximum inversion (MAX_inversion_) angles during stance phase, and ankle frontal angle at initial contact (IC_inversion_). Shoe stack heights were abbreviated as H: 50 mm, M: 35 mm & L: 27 mm. Descriptive statistics is given as mean ± standard deviation. The *post-hoc* tests were calculated for mean data at 10 km/h and 15 km/h. Significant results are highlighted in bold.

| $t_{\text{eversion}}$ (%) | | | | | | | | |
| --- | --- | --- | --- | --- | --- | --- | --- | --- |
| | 10 km/h | 15 km/h | rmANOVA | p | $\eta^2$ | Post-hoc | p | \|d\| |
| H | 21.5 ± 27.2 | 24.8 ± 27.2 | Shoe | **0.035** | **0.189** | H vs. M | **0.006** | **0.83** |
| M | 13.2 ± 21.7 | 14.2 ± 22.1 | Speed | 0.504 | 0.028 | H vs. L | 0.371 | 0.08 |
| L | 22.1 ± 26.7 | 21.2 ± 21.0 | Interaction | 0.423 | 0.052 | M vs. L | **0.046** | **0.52** |
| $\text{MAX}_{\text{eversion}}$ (°) | | | | | | | | |
| | 10 km/h | 15 km/h | rmANOVA | p | $\eta^2$ | Post-hoc | p | \|d\| |
| H | 1.0 ± 4.7 | 1.6 ± 6.0 | Shoe | **0.006** | **0.271** | H vs. M | **0.002** | **0.92** |
| M | -1.3 ± 4.0 | -1.1 ± 4.5 | Speed | 0.332 | 0.059 | H vs. L | 0.388 | 0.07 |
| L | 0.8 ± 4.1 | 1.2 ± 3.7 | Interaction | 0.859 | 0.009 | M vs. L | **0.010** | **0.71** |
| $\text{MAX}_{\text{inversion}}$ (°) | | | | | | | | |
| | 10 km/h | 15 km/h | rmANOVA | p | $\eta^2$ | Post-hoc | p | \|d\| |
| H | -10.9 ± 4.5 | -12.9 ± 5.5 | Shoe | **0.005** | **0.284** | H vs. M | **0.002** | **0.90** |
| M | -12.5 ± 4.8 | -13.8 ± 5.3 | Speed | **<0.001** | **0.555** | H vs. L | 0.430 | 0.04 |
| L | -10.8 ± 3.5 | -12.9 ± 4.9 | Interaction | 0.583 | 0.033 | M vs. L | **0.004** | **0.85** |
| $\text{IC}_{\text{inversion}}$ (°) | | | | | | | | |
| | 10 km/h | 15 km/h | rmANOVA | p | $\eta^2$ | | | |
| H | -10.3 ± 4.6 | -11.7 ± 6.0 | Shoe | 0.057 | 0.192 | | | |
| M | -11.8 ± 5.4 | -12.8 ± 5.6 | Speed | **0.001** | **0.480** | | | |
| L | -8.8 ± 6.2 | -11.3 ± 5.7 | Interaction | 0.477 | 0.045 | | | |

#### 3.2.2 Nonlinear analysis

The results are summarized in Table 4. The λ_hip_ showed significant effects for the factor shoe (p = 0.040). The *post-hoc* results showed a reduced local stability in the hip for High compared to Low (p = 0.027). Head, trunk and foot did not show significant effects.

**Table 4:** The parameters used in the nonlinear analysis of running stability: local dynamic stability of head (λ_head_), trunk (λ_trunk_), hip (λ_hip_) & foot (λ_foot_); detrended fluctuation scaling component of stride time (DFA-α). Shoe stack heights were abbreviated as H: 50 mm, M: 35 mm & L: 27 mm. Descriptive statistics is given as mean ± standard deviation. The *post-hoc* tests were calculated for mean data at 10 km/h and 15 km/h. Significant results are highlighted in bold.

| $\lambda_{\text{head}}$ | | | | | | | | |
| --- | --- | --- | --- | --- | --- | --- | --- | --- |
| | 10 km/h | 15 km/h | rmANOVA | p | $\eta^2$ | | | |
| H | 1.52 ± 0.12 | 1.52 ± 0.14 | Shoe | 0.423 | 0.052 |  |  |  |
| M | 1.53 ± 0.16 | 1.51 ± 0.11 | Speed | 0.605 | 0.017 |  |  |  |
| L | 1.50 ± 0.14 | 1.48 ± 0.11 | Interaction | 0.903 | 0.006 |  |  |  |
| $\lambda_{\text{trunk}}$ | | | | | | | | |
| | 10 km/h | 15 km/h | rmANOVA | p | $\eta^2$ | | | |
| H | 1.11 ± 0.07 | 1.10 ± 0.13 | Shoe | 0.233 | 0.859 |  |  |  |
| M | 1.12 ± 0.14 | 1.12 ± 0.08 | Speed | 0.123 | 0.201 |  |  |  |
| L | 1.11 ± 0.10 | 1.04 ± 0.12 | Interaction | 0.124 | 0.610 |  |  |  |
| $\lambda_{\text{hip}}$ | | | | | | | | |
| | 10 km/h | 15 km/h | rmANOVA | p | $\eta^2$ | *Post-hoc* | p | \|d\| |
| H | 1.48 ± 0.13 | 1.43 ± 0.15 | Shoe | **0.040** | **0.182** | H vs. M | 0.241 | 0.17 |
| M | 1.46 ± 0.16 | 1.41 ± 0.14 | Speed | 0.253 | 0.081 | H vs. L | **0.027** | **0.64** |
| L | 1.38 ± 0.14 | 1.39 ± 0.01 | Interaction | 0.373 | 0.060 | M vs. L | 0.115 | 0.40 |
| $\lambda_{\text{foot}}$ | | | | | | | | |
| | 10 km/h | 15 km/h | rmANOVA | p | $\eta^2$ | | | |
| H | 1.05 ± 0.12 | 1.06 ± 0.10 | Shoe | 0.859 | 0.009 |  |  |  |
| M | 1.01 ± 0.16 | 1.08 ± 0.13 | Speed | 0.201 | 0.100 |  |  |  |
| L | 1.03 ± 0.13 | 1.08 ± 0.16 | Interaction | 0.610 | 0.030 |  |  |  |
| DFA-α | | | | | | | | |
|  | 10 km/h | 15 km/h | rmANOVA | p | eta |  |  |  |
| H | 0.66 ± 0.17 | 0.83 ± 0.21 | Shoe | 0.127 | 0.121 |  |  |  |
| M | 0.66 ± 0.23 | 0.91 ± 0.20 | Speed | **<0.001** | **0.725** |  |  |  |
| L | 0.58 ± 0.18 | 0.83 ± 0.24 | Interaction | 0.505 | 0.042 |  |  |  |

## 4 Discussion

This study aimed to understand the effects of different shoe stack heights on running style and stability during treadmill running at two different speeds. The results showed that the higher stack heights can affect running style (i.e. SF_norm_, DF, COM_osc_, k_ver_, sagittal and frontal ankle angles) and reduce stability (i.e. t_eversion_ MAX_eversion_, MAX_inversion_ and **λ_hip_**), in line with our first and second hypotheses. More concretely, High resulted in longer ground contacts relative to the stride time compared to Low. Furthermore, the higher the stack height, the lower was the COM_osc_, whereas the k_ver_ was the highest for Low with no differences between Medium and High. The ankle was longer in an everted position with High compared to Medium. In addition, the local dynamic stability of the hip during running was lower with High compared to Low. The higher stack heights (≥ 35 mm) led to a lower step frequency normalized to leg length at 15 km/h but not at 10 km/h. Except for SF_norm_, the findings were not speed dependent but global effects, which contradicts to our third hypothesis.

### 4.1 Higher shoe stack height leads to changes in running style parameters that are associated with a decreased running economy

Running style is important for economy and performance (Folland et al., 2017). It was hypothesized here that different shoe stack heights modulate the running style (H1). This study revealed that, at a greater speed, the higher stack heights (High and Medium) led to a lower SF_norm_ compared to the lowest stack height. This can be interpreted as a shift of the running style to the direction of the “push” style associated with longer steps (van Oeveren et al., 2021). In addition, independent of running speed, the highest stack height led to higher DF compared with the lowest stack height, which can be interpreted as a running style shifted to the direction of a “stick” style associated with longer stance time relative to stride time (van Oeveren et al., 2021). When it comes to the ratio of braking to propulsion durations (i.e. L2T), the results showed that the braking phase had a shorter duration than the propulsion, but L2T remained unaffected by different shoes and running speeds. To sum up, the highest stack height altered the running style in line with our first hypothesis and led to longer steps (based on SF_norm_) with longer ground contacts relative to the stride time (based on DF), especially compared to the lowest stack height and faster speed. However, it should be added that the differences between shoe conditions were 2-4% for SF_norm_ and 1.5% for DF. This study does not address whether these changes can translate into performance differences.

Further analysis revealed that COM_osc_ increased with increasing stack height. This finding may indicate a decreased running economy for the highest stack height condition, since a larger COM_osc_ is associated with worse running economy (Folland et al., 2017; Saunders et al., 2004). Nevertheless, it should be noted that running economy was not directly measured therefore this is not necessarily true. For example, Fletcher et al. (2008) detected differences in COM_osc_ but not in running economy. Therefore, this interpretation should be treated with caution. Furthermore, k_ver_ was the highest for Low with no differences between Medium and Heigh.

The results of k_leg_ showed significant speed effects but no shoe effects. This secondary finding is surprising since k_leg_ takes effective leg length into account, thereby eliminating the speed dependency; unlike k_ver_ which increases with running speed. Normally, it is expected that k_leg_ remains relatively stable across different running speeds (Brughelli & Cronin, 2008) which was not the case in the current study. Possibly, it is because the stiffness parameters were estimated based on kinematics without any force measurements, therefore possibly they are less reliable compared to measures based on kinetics. However, the kinematics-based method has also been used in various other studies to estimate stiffness (e.g., Burns et al., 2021; Morin et al., 2005).

Finally, the analysis of joint angle time series of the lower limb showed that the different stack heights affect joint kinematics to a certain extent. In the frontal plane, the middle stack height shoes led to a decreased frontal ankle angle (shift to inversion) compared to other shoes, whereas in the sagittal plane there were no significant effects in the *post-hoc* tests. These results indicated that the different shoe stack heights affect mainly the ankle but not the more proximal joints of the lower body (i.e. the knee and hip).

### 4.2 Reduced running stability with highest stack height

In this study, it was hypothesized that the shoes with higher stack heights lead to less stability (H2), in line with previous findings (Hoogkamer, 2020; Ruiz-Alias, Jaén-Carrillo, et al., 2023). Both linear and nonlinear analyses were carried out in a comprehensive approach. Linear measures provide essential information such as mean and standard deviation on the analyzed parameters but they assume that the output of the system is directly proportional to the input although biological systems are inherently nonlinear (McCamley & Harrison, 2016). Therefore, nonlinear analysis complements linear analysis by providing deeper insights to biological systems without simple linearity assumptions. The linear analysis of joint angles based on discrete parameters revealed that Medium led to a lower MAX_inversion_ and MAX_eversion_ during stance. It is important to note that MAX_eversion_ values were partially still in the region of an inversion (Fig. 4). Together with the joint angle time series data, it can be concluded that Medium shifted the frontal ankle angle to a more inverted position compared to High and Low. The t_eversion_ results revealed that High led to a longer duration in an everted foot position compared to Medium, which may be indicator for a decreased stability and a higher risk of injury (Becker et al., 2018; Hannigan & Pollard, 2020). However, it should be noted that the shoes with the highest stack were not less stable compared to the lowest stack height shoes. This may be attributed to the missing advanced footwear technology elements in Low. Possibly, the advanced footwear technology elements (i.e. curved carbon infused rods) help to increase ankle stability but extremely high stack heights (i.e. 50 mm) counterbalance the advantages of these technologies. Thereby, the linear analysis results partially supported our second hypothesis. These findings were in line with those from Barrons et al. (2023a) but not fully matched them. In their study, the shoes with 35 mm stack height decreased ankle eversion compared with 45 mm but not with 50 mm. It should also be noted that the frontal ankle angle values were slightly different from those in this study, even though the tendencies due to different shoes were similar. These differences were probably due to different marker sets and inverse kinematics models used. Previous studies reported that the kinematics results may change, especially in the non-sagittal planes, when different setups are used for gait analysis (Roelker et al., 2017; Trinler et al., 2019; Wouda et al., 2018).

The nonlinear analysis indicated that local stability of the hip based on MLE was lower with High compared to Low. In this context, hip stability can be seen as the most important location since the fundamental goal of locomotion is to transport the COM (Evans et al., 2022) and the hip can be used to approximate the COM during running (Napier et al., 2020). The MLE results indicated that stabilization of the hip became more difficult with the highest stack height compared to the lowest, which thereby supported our second hypothesis. The DFA of stride time indicated a self-similarity characteristic (e.g., a short stride is more likely to follow a short stride). In contrast to MLE, the DFA of stride time did not show any significant shoe differences, which was against our second hypothesis. One explanation may be that the shoes provide only small perturbations for the system (Prejean & Ricard, 2019). Therefore, they do not modulate the global stability.

In this study, one of the goals was to operationalize stability from different perspectives to gain a better understanding of stack height effects on stability. First, the discrete frontal ankle angles indicated a higher portion of eversion during stance for High compared to Medium. Secondly, the local dynamic stability of the hip during running was lower with High compared to Low. These results are in line with the perceived stability results of the data used in this study (Fadillioglu et al., 2024) as well as with previous studies suggesting that increasing stack height may lead to instabilities (Hoogkamer, 2020; Ruiz-Alias et al., 2023a). Based on these findings, it can be suggested that high stack height (i.e. 50 mm) lead to instabilities in frontal ankle angle and hip local stability. But it is important to note that High and Medium had curved carbon infused rods but Low did not. The detected differences may partially be attributed to the carbon rods and not solely to the higher stack heights. It should be added that High had a stack height larger than the allowed limit (40 mm) by World Athletics regulation (World Athletic Council, 2022). On this basis, in terms of the stability concerns, the findings of this study support this regulation restricting the stack height to 40 mm. However, the current study does not address whether the observed decreased instability can be translated into a decreased running performance. Therefore, further research is needed. Furthermore, it should be noted that not all parameters supported this statement. For example, the local dynamic stability of the foot did not change. Nevertheless, this is not in conflict with the frontal ankle angle results since linear and nonlinear analyses do not have the same assumptions (McCamley & Harrison, 2016).

### 4.3 Stack height changes were largely independent of running speed

In general, higher running speeds may represent more challenging task conditions than lower speeds (Santuz et al. 2020b), thereby increase the influence of stack heights. In this study, it was hypothesized that the effects of different stack heights would be pronounced at higher speeds (H3).

The detected shoe effects did not change between the two running speeds, and only SF_norm_ had significant interaction effect between the factors speed and shoe. More concretely, the shoe effects were visible at a higher speed only for SF_norm_. Based on these findings, it can be suggested that the detected shoe effects were mostly global effects, which means that they were independent of the tested running speeds, with the exception of the running style parameter SF_norm_. Therefore, our third hypothesis should be rejected with the exception of the SF_norm_.

### 4.4 Limitations

This study has some limitations that must be mentioned. Firstly, the shoes differed mainly in their stack heights but not all the remaining shoe features were identical. Major limitation was that the shoes with the highest and middle high stack height included advanced footwear technology elements (carbon curved rods) but the lowest stack height shoes did not contain these elements. The changes between High and Medium can mainly be attributed to stack height changes whereas the changes compared to Low can be additionally due to infused carbon rods in the shoe soles (e.g., local stability changes of the hip). Furthermore, the weight of the shoes slightly differed but the difference between the lightest and heaviest ones was only 49 g (Table 1). It has previously been shown that an added mass of 50 g resulted in limited biomechanical changes (Rodrigo-Carranza et al., 2020). Secondly, the experiments were done on a treadmill at two constant running speeds. Even if it is done in a similar way in many comparable studies also (e.g., Barrons et al., 2023a; Chambon et al., 2014; Udin et al., 2023), it should be kept in mind that this is a very controlled experimental condition. It is possible that larger additional perturbations to the system (e.g., higher running speed, uneven ground and fatigue) will increase the shoe effects. Thirdly, the participants were healthy experienced runners and the findings cannot be directly translated to other expertise levels (Fadillioglu et al., 2022; Möhler et al., 2020). Fourthly, the initial contact and toe-off events were detected based on kinematics in the current study. Even if the validation studies reported the difference between gold standard and proposed methods to be low for treadmill running (error ≤ 20 ms (Fellin et al., 2010; King et al., 2019; Leitch et al., 2010)), the kinetic-based methods are more accurate. Lastly, the used dual-axis framework (van Oeveren et al., 2021) was proposed to provide an overview of running style with fundamental differences between conditions. However, this framework is not enough to describe a running style completely but proper normalization for individual characteristics is required.

## 5 Conclusion

The purpose of this study was to investigate how three shoes with different stack heights affect running style and stability during treadmill running at different speeds. The key findings of this study were: (1) changes in stack height can affect running style. Particularly, the shoes with the highest stack height resulted in longer ground contacts relative to the stride time compared to the lowest stack. Furthermore, the higher the stack height, the higher was the vertical oscillation of the COM (2) The ankle spent longer in an everted position with the highest stack compared to the middle heigh. In addition, the local dynamic stability of the hip during running was lower with the highest stack compared to the lowest one. Both results indicated that the highest stack height (50 mm) reduce stability. However, not all stability parameters indicated decreased stability for the highest stack height. (3) The higher stack heights (≥ 35 mm) led to a lower step frequency normalized to leg length at 15 km/h but not at 10 km/h. The remaining shoe effects were independent of the running speed.

Shoe stack height is a highly discussed topic especially since the introduction of a stack height regulation by World Athletics. The findings of the study are in line with the statement “more is not always better” when it comes to the stack height. This study has found another piece of puzzle in understanding the effects of stack heights on running style and stability. Future studies may focus on running coordination as well as analysis of joint loads and muscle activities to better understand stack height effects.

## Acknowledgements

We acknowledge support by the KIT-Publication Fund of the Karlsruhe Institute of Technology.

## Ethics statements

### Studies involving animal subjects

No animal studies are presented in this manuscript.

### Studies involving human subjects

The studies involving humans were approved by the ethics committee of the Karlsruhe Institute of Technology (KIT). The studies were conducted in accordance with the local legislation and institutional requirements. The participants provided their written informed consent to participate in this study.

### Inclusion of identifiable human data

No potentially identifiable images or data are presented in this study.

## Data availability statement

The raw data supporting the conclusions of this article will be made available by the authors, without undue reservation.

## Generative AI disclosure

No Generative AI was used in the preparation of this manuscript.

